# Impacts of the 1918 flu on survivors’ nutritional status: a double quasi-natural experiment

**DOI:** 10.1101/2020.04.23.057638

**Authors:** Alberto Palloni, Mary McEniry, Yiyue Huangfu, Hiram Beltran-Sanchez

## Abstract

A unique set of events that took place in Puerto Rico during 1918-1919 generated conditions of a “double “quasi-natural experiment. We exploit these conditions to empirically identify effects of exposure to the 1918 flu pandemic, those of the devastation left by an earthquake-tsunami that struck the island in 1918, and those associated with the joint occurrence of these events. We use geographic variation to identify the effects of the quake and timing of birth variation to identify those of the flu. In addition, we use markers of nutritional status gathered in a nationally representative sample of individuals aged 75 and older in 2002. This unique data set enables to make two distinct contributions. First, unlike most fetal-origins research that singles out early nutritional status as a *determinant of adult health*, we test the hypothesis that the 1918 flu had deleterious effects on the nutritional status on adult survivors who at the time of the flu were *in utero* or infants. Second, and unlike most research on the effects of the flu, we focus on markers of nutritional status set when the adult survivors were children or adolescents. We find that estimates of effects of the pandemic are sizeable primarily among females and among those who, in addition to the flu, *were exposed to the earthquake-tsunami.* We argue that these findings constitute empirical evidence supporting the conjecture that effects of the 1918 flu alone and the combined effects of the flu and the earthquake are associated not just with damage experienced during the fetal period but also postnatally.

## INTRODUCTION

The Spanish flu virus of 1918-19 is an example of a perfect storm: like HIV and unlike ordinary standard seasonal influenza, it was highly lethal but, unlike HIV and like other influenza, it was rapidly and efficiently spread ^(1-5)^. The combination of these two traits made the pandemic one of the deadliest in human history ^(6, 7)^. In most of the world the A/H1N1 influenza was characterized by its unusual temporal sequence, peculiar age pattern, and case morbidity and lethality ^(1, 8, 9)^. The manifestation of the pandemic in Puerto Rico followed the world pattern closely but, as we will see later, it added unique features. Jointly, the age pattern of incidence and morbidity and mortality levels, created unfavorable conditions for all but especially for women of childbearing ages and those who were pregnant at the time or who had recently given birth ^(10, 11)^. These conditions may have compromised not just fetal growth but also infant and young children’ health, both highly dependent on maternal health status and parental care.

As if the onslaught of the flu had not been enough, on October of 1918, precisely when the pandemic was gathering force during its second, most lethal wave, a strong earthquake (the San Fermin earthquake) struck the Western part of the island. This was immediately followed by a tsunami, two major aftershocks in a two-month interval following the earthquake, and multiple smaller ones spread over the subsequent year or so ^(12)^

A large body of research on the lasting effects of the 1918 pandemic relies on the fact that the event can be considered a quasi-experiment: it was unexpected, difficult to avoid and, in most cases, there were no contemporaneous exogenous events that could have produced similar outcomes^(1, 10)^. The Puerto Rican earthquake was also unexpected, hard to avoid in areas struck by it and, asides from the flu pandemic, unaccompanied by other major events that could have injured with equal violence an already vulnerable population. Thus, in one stroke, an unlikely combination of two events handed us conditions of a unique double quasi experiment.

This paper departs form others on effects of 1918 flu pandemic. First, it seeks to shed light on a rather unexplored dimension of the 1918 pandemic, namely, its effects on *markers of nutritional status* of individuals exposed to it. With the exception of one study ^(10)^, we know of no other attempt to investigate such an association. Analyses of impacts on the nutritional status of 1918 flu survivors requires to focus on mechanisms that could disturb physiological growth and developmental processes during infancy, early childhood and even early adolescence, not just those that operate *in utero*. It is known that embryonic and, more generally, intrauterine disruptions influence neural development (brain tissue), metabolic balance (pancreas, liver), nephron growth (kidneys and regulation of blood pressure) or lung and heart functioning ^(13-16)^. In addition, embryonic and fetal development is also about growth of cartilage, bone and muscle tissue, all of which are implicated in subsequent postnatal physical development ^(17)^. In addition, impairment of growth processes that occur during the fetal period can be aggravated if postnatal conditions deteriorate. Thus, fetal growth could be impaired when pregnant mothers experience illnesses and are exposed to either episodic or chronic stress. By the same token, when due to illnesses or death, mothers cannot breastfeed normally, are unable to provide sufficient maternal care, proper nutrition, grooming, and hygiene, early growth and development could go astray. Furthermore, when infants and young children face prolonged exposure to adverse environmental and material conditions, catch-up growth may be a non-starter ^(18, 19)^. This justifies the need to assess not just the 1918 flu ‘s impacts of in *in-utero* exposure, but also those closely associated with adverse postnatal conditions.

Second, we build the case on a unique quasi-experimental research design, a product of the occurrence of two simultaneous events, one involving *timing of exposure* (flu) and the other involving *timing and geography of exposure* (flu and regional earthquake-tsunami). We aim to show that the flu pandemic and the earthquake-tsunami combine to generate impacts that neither of these events could have produced separately and are strongly associated with both gestational and postnatal exposures.

### Early physical growth debilitation and its long run consequences

Human physical growth depends on early embryonic and fetal events, maternal exposures (including stressors), maternal health status, and parental effects, including maternal capacity to nourish during the fetal and postnatal stages ^(20-22)^.Of particular importance is the length and intensity of breastfeeding ^(23-25)^, protection from infections and parasitic diseases ^(26)^, recovery from illness^(27, 28)^ and reduction of environmental stressors^(29)^. These parental effects are strongly associated with maternal (and paternal) health status, household (family) environments and access to resources.

#### Embryonic and fetal growth

By and large, fetal nutrition depends on maternal diet and placental capacity to deliver nutrients (including oxygen, fat, proteins, hormones, SCFA)^(30)^. It is well-known that maternal nutritional status influences the entire process of fetal development and can have strong impacts of the infant’s subsequent growth^(31)^. It is also known that poor maternal health status can derail the normal course of a pregnancy and complicate delivery. In particular, maternal infections during pregnancy could compromise normal fetal development and their ultimate impacts depend on the timing of infections, their intensity, and duration. These effects are also associated with inflammatory responses triggered by the infections. In addition to the potentially fetal organogenic damage associated with the flu-related cytokine storms ^(2, 32)^, bouts of maternal hyperthemia induced by inflammation can also lead to deleterious outcomes, including miscarriages, premature labor, stillbirths, congenital anomalies, and growth restrictions ^(33-35)^. The latter are a result of irregularities of the physiology of bone and muscle tissues formation as bone develops from embryonic mesoderm and proceeds by ossification of cartilage tissue formed from mesenchyme. Maternal hyperthermia can also affect limb myogenesis as it disrupts and delays the involvement of several crucial regulatory factors^(35)^. Jointly, dysregulation of bone and muscle tissue formation can compromise normal physical growth ^(36)^.

#### Early and late infant development

Because of mother’s milk properties, intensity and length of breastfeeding are of crucial importance for infants’ early growth, particularly during the first 6 months of life ^(37)^. Aside from its beneficial nutritional properties ^(38)^, breastmilk contains important compounds that strengthen infants’ immune response and act as a shield to reduce risks of disease^(39)^. Most viral, bacterial and parasitic diseases reduce appetite, limit food intake and impair the child’s nutrient absorption capabilities ^(40)^. Thus, the combination of illnesses and breastfeeding interruption, cessation, or irregularities during the first 6 months can compromise not only the quality and quantity of nutrients available for early growth but also reduce absorption and metabolization of those available ^(41)^. These disruptions compromise the ability of an organism to satisfy energetic demands to sustain rapid cell division and specialization and organ growth and formation during critical periods ^(30)^. Although early growth faltering can be offset by subsequent catch-up growth phases, this will not take place in the absence of material conditions that can sustain rapid growth and maturation ^(19, 42)^. In populations with widespread poverty and vulnerable maternal health status, the process of catch-up growth may never get off the ground and children who could have benefitted from it will fail to attain physical growth milestones^(42)^.

### Long lasting effects of the flu

These considerations lead us to hypothesize that exposure to the flu during critical periods as we define them here, e.g in utero and/or during infancy, must have had non-negligible influences on early nutritional status and should be reflected in poor adult markers of physical growth. By the same token, exposure to stresses and material deprivation brought about by the earthquake-tsunami could have disrupted embryonic, fetal and postnatal growth and, as consequence, facilitated growth faltering and attainment of substandard markers of physical growth. Furthermore, as did happen in other populations, the flu effects were probably stronger among those who experienced the pandemic in areas more severely affected by it ^(1, 10)^. Finally, both *in utero* and postnatal vulnerability to the flu was likely augmented by conditions associated with the earthquake^(43)^. If so, we should find that the impact of the flu among the “treated” by the flu (e.g. those exposed to the pandemic *in utero* or during the first year of life) and “controls” (e.g. those exposed later in childhood or adolescence) is larger among those born in areas struck by the earthquake-tsunami (e.g. “treated” by the earthquake) than among those born elsewhere in the island (“controls”). Table 1 is a stylized representation of the study design.

**Table 1:** Stylized representation of study design.

| Table 1: Stylized representation of study design |  |  |
| --- | --- | --- |
|  | **Early Life Earthquake Exposure** |  |
| **Early Life Flu Exposure** | Yes (“treated”) | No (“not treated”) |
| Yes (“treated”) |  |  |
| Born 1918-1919 | A | B |
| No (“controls”) |  |  |
| Born before 1917 | C | D |
| Born after 1920 | E | F |
| Notes: The key contrasts we examine in the paper are: |  |  |
| (i) | A vs B; (C+E) vs (D+F) | effects of earthquake |
| (ii) | A vs (C+E) | effects of flu among those not born in earthquake areas |
| (iii) | A vs (D+F) | effects of flu among those born in earthquake areas |
| (iv) | A vs (B+C+D+E+F) | gross effects of the flu |

An ancillary issue relates to potential gender differentials of the flu effects. Although there are behavioral mechanisms that could have exacerbated impacts among female infants (e.g male children preferences), gender differences may surface as a result of culling among males. Because male embryos are more vulnerable ^(17)^ and male infants experience higher mortality than female infants^(44)^, one may observe stronger effects among females than among males as a result of selection.

## MATERIALS AND METHODS

### Data

We use PREHCO (Puerto Rican Elderly Health Conditions) data base^(45)^. PREHCO is a two-wave panel of the non-institutionalized Puerto Rican population aged 60 and over and their surviving spouses. The study uses a multistage, stratified sample of the elderly population residing in Puerto Rico in the year 2002 with oversamples of regions heavily populated by people of African descent and of individuals aged over 80. A total of 4,293 in-home face-to-face target interviews were conducted between May 2002 and May 2003 and a second wave data were collected during 2006-2007. The overall response rate was 93.9%. Our analyses use a subpopulation aged 74+ at the time of first interview, e.g. those born between 1896 and 1927. The total sample size is 1,613 observations, 956 of them females. About 30 percent of the sample were born on or before 1917 and 11 percent between years 1918 and 1919. A histogram of the distribution of year of birth is in Fig 1 and a summary of key statistics is in Table 2.

**Table 2:**
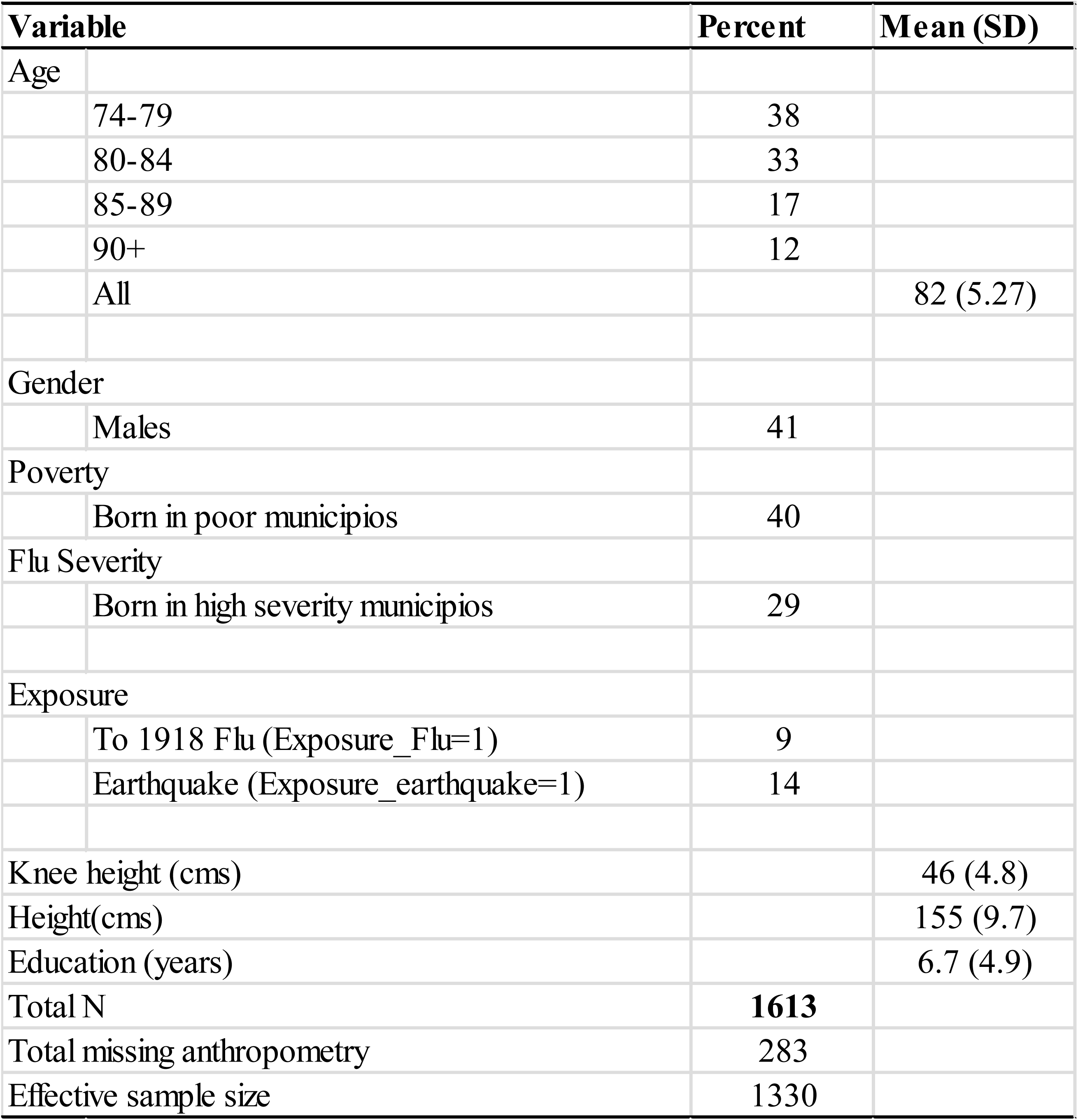
Summary of selected sample statistics.

**Fig 1:**
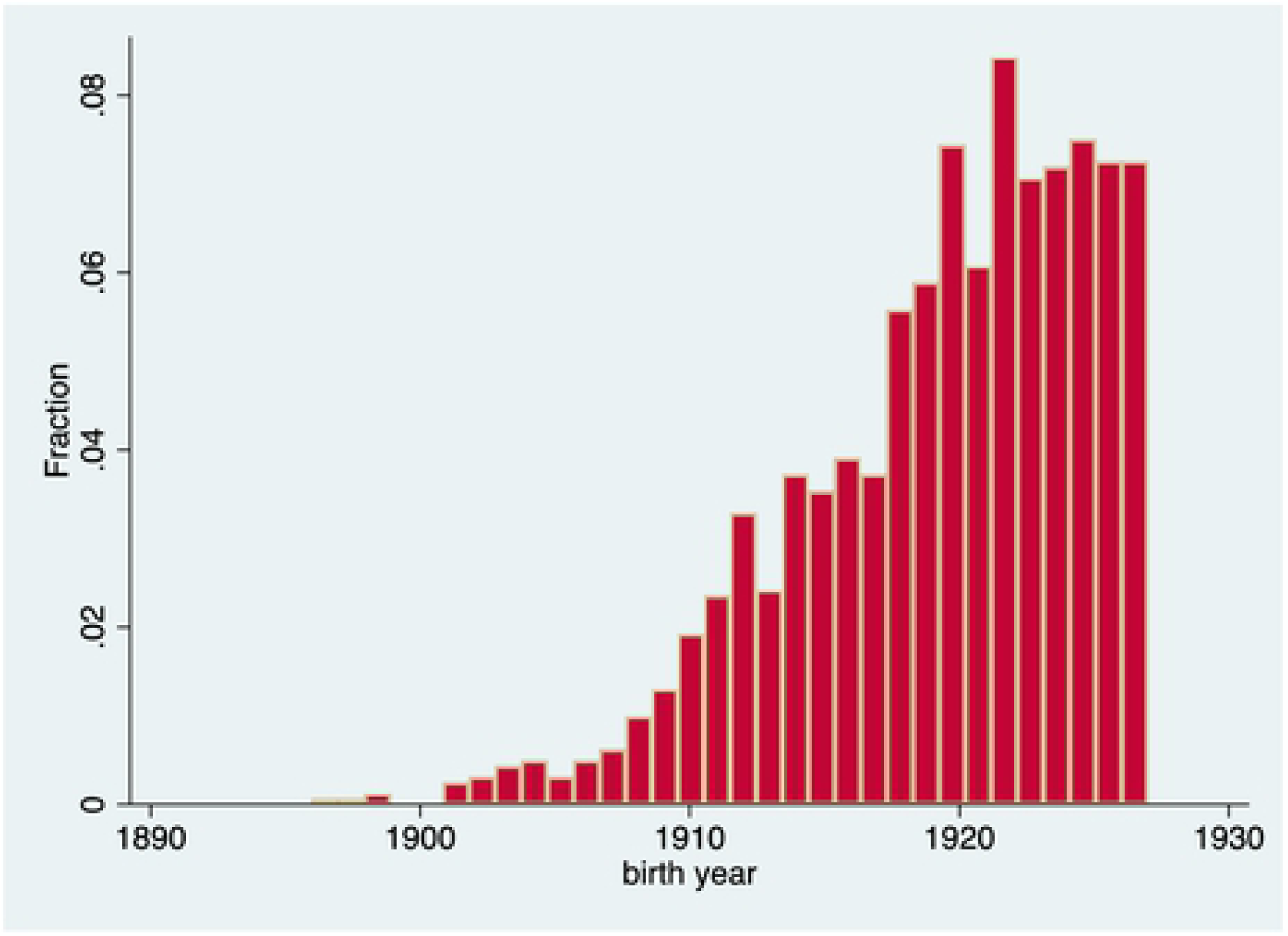
Distribution of year of birth.

### Measures

#### Flu exposure

In contrast to other studies of the 1918 flu, we identify a wider period during which exposure is assumed to have taken place and, in addition to the fetal period, we also include time intervals during which post-natal care may have been disrupted as a consequence of the epidemic and/or the earthquake. Note that if, as we argue here, the post-natal mechanisms are also relevant for outcomes other than physical growth markers (adult health, mortality, cognition, educational attainment, etc…), studies that ignore them will underestimate the total effects. This is because when a restrictive definition of exposure is used some “treated” cases will be assumed to be “controls”. To capture extended exposure including gestational and post-natal exposures we define a dummy variable attaining the value 1 if an individual’s birth is reported to have taken place during 1918 or during the first six months of 1919. This indicator is a compromise between preservation of the ability to assign effects to *fetal and post-natal exposure according to timing and duration*, on one hand, and sample size constraints, on the other. A full rationale for this choice is provided in a lengthy description of the association between year and month of birth and type and degree of exposure to the various waves of the 1918 pandemic and earthquake. *(reference omitted to preserve anonymity)*

#### Flu Severity

Although we lack information on the incidence and case fatality of the pandemic in Puerto Rico, we follow past research and create a proxy indicator of flu severity using the excess total mortality registered during the flu period ^(1)^. To construct an index of severity we consider total mortality during the two years period 1918-1919 for each of the 76 municipios (the smallest administrative units) in Puerto Rico, estimate expected deaths using age-specific mortality rates during 1918 in the US, and then compute the ratio of mortality rate observed in a municipio to the observed rate. Note that this quantity is equivalent to an indirectly standardized mortality ratio, a conventional index computed when information of age specific death rates is absent. The information on municipio’s mortality is retrieved from Luk’s estimates ^(46)^.

We classify as high severity all municipios above the 90^th^ centile of the severity index distribution (details of the index construction are in S1 Text).

#### Poverty

We adopt the classification of municipios constructed by Clark ^(47)^. Municipalities were grouped into three classes according to their population size, assessed value, and government income. A total of 25 municipalities are either in the wealthiest or an intermediate class and the remaining municipalities are in the poorest category. In this paper we use a 0/1 binary indicator to contrast the poorest and the remaining municipios.

#### Earthquake-tsunami exposure

Exposure to earthquake-tsunami is assessed according to municipio of birth. We classify these into three groups depending on the severity of the event: (i) most severe, (ii) severe and (iii) not severe. In what follows we use a 0/1 dummy variable to flag municipios in group (i). Group (i) includes the municipios of Aguada, Aguadilla, Anasco, Isabella and Mayaguez. Group (ii) includes the rest of the West Coast municipios (Cabo Rojo, Hormigueros, Rincon, San Sebastian and Quebradilla). The remaining municipios are in group (iii). This grouping is based on historical accounts of the earthquake-tsunami and is consistent with the geographic location of municipios relative to the epicenter of the earthquake and exposure to the tsunami that accompanied it.

#### Knee height and adjusted height

We use PREHCO’s anthropometry module for the assessment of height and knee height. To attenuate biases due to skeletal compression, we adjust height measures using estimates of compression by gender and age observed in a sample of individuals who were followed for a long period of time (see S2 Text). The magnitude of the adjustments is considerable and, if anything, they will lead to overcorrections and to downwards biases of the effects on height of exposure to events of interest. To circumvent the problem altogether we also use knee height, a marker of early nutritional status unaffected by skeletal compression. There are a number of outcomes frequently studied in the literature on the 1918 pandemic, including BMI. We do not examine these since our interest is on markers of *early* nutritional status and neither BMI nor any of other available in the survey are suitable.

### Models

We use seemingly unrelated regression (SUR) and treat adjusted height and knee height as continuous variables with possibly correlated errors. We estimated three alternative classes of models, including SUR, OLS and bivariate probit. Although they all lead to the same inferences, we only discuss results associated with SUR models because they produce easily interpretable estimates, do not depend on arbitrary cut points (as bivariate probit models do), and generate more conservative standard errors than OLS.

The SUR model contains two equations, one for each continuous trait, with potentially different covariates in each and assumed correlated errors. While the estimates of *separate* OLS equations are consistent the estimated standard errors are inconsistent and possibly subestimated. In all cases we use our preferred measures of exposures, namely, *Exposure_flu* for flu and *Earthquake Exposure* for earthquake-tsunami exposure The specification for outcome j=1(knee height) and j=2 (adjusted height) is

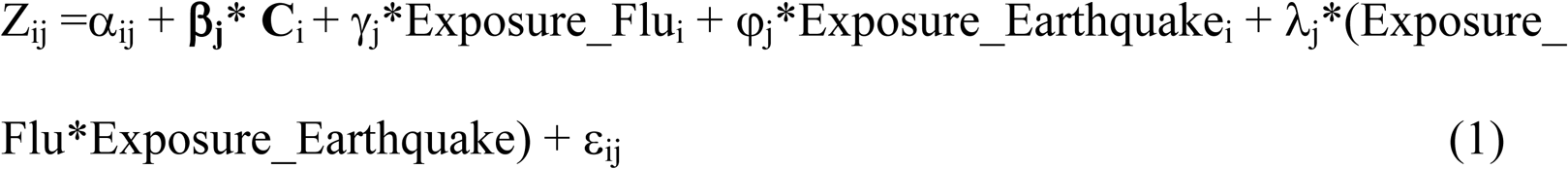

where Z_ij_ is the outcome of interest for individual i, **C**_i_ is a vector of control variables for individual i, *Exposure_Flu* is a 0/1 variable for exposure to flu, *Exposure_Earthquake* is a 0/1 variable for exposure to earthquake-tsunami, and ε_ij_ is an idiosyncratic error. In turn, the parameters for the equation of outcome j are a constant, α_j_, a vector of effects associated with controls, **β**_**j**_, the effect of flu exposure, γ_j_, the effect of earthquake-tsunami exposure, φ_j_, and the effect of interaction terms, λ_j_. The vector of control variables includes years of education (continuous) and municipio of birth’ s poverty level (discrete). Additional controls were discarded since they did not change results.

Three remarks are important. First, our model specification *does not* include a control for age as there is no relation between markers of nutritional status and age. Second, the regression formulation is a difference-in-difference (DiD) model that seeks to identify (i) differences in the impact of the flu by timing of exposure between high severity and low severity areas and (ii) differences in the impact of the flu by timing of exposure in geographic areas affected by the earthquake-tsunami (high severity of earthquake) and those not affected (low severity area). Unlike standard DiD models, we are estimating differences across two, not just one “treatment”. Finally, to minimize impacts of culling, all models were estimated separately for males and females.

## RESULTS

### Baseline models

Table 3 shows that it is only among females that exposure to flu and to earthquake have noticeable effects on both knee height and adjusted height. These effects are quite large, properly signed, and associated with p-values less than [.01,.02], except for effects of earthquake exposure on adjusted height. The reduction in knee height implied by the estimated effect of flu exposure is about .33 of a standard deviation of knee height’s and equivalent to .033 of the observed mean (see Table 2). The corresponding quantities for exposure to earthquake are .23 and .022 respectively. The *relative* impacts on height are slightly smaller. As shown below, estimates of effects for females are always large, systematic, robust, and lead to unequivocal inferences. In contrast, estimates for males tend to be of smaller magnitude and less systematic. For this reason, we only discuss estimates for females. It should be remembered that this contrast was anticipated as part of the conjecture that males experience considerably more culling that females. (To avoid cluttering, results for males are displayed in Table in S3 Table).

**Table 3:** Baseline models by gender.

| Table 3: Baseline models by gender |  |  |  |  |
| --- | --- | --- | --- | --- |
|  | Females |  | Males |  |
|  | (1) | (2) | (3) | (4) |
| Variables | Knee | Adj. height | Knee | Adj. height |
| Years Education | 0.051<br>[1.599] | 0.290<br>[5.842] | 0.101<br>[2.482] | 0.239<br>[3.569] |
| Exposure Flu (1/0) | -1.082<br>[-2.048] | -2.289<br>[-2.771] | -1.016<br>[-1.425] | 0.492<br>[0.421] |
| Poverty Birthplace (1/0) | -0.628<br>[-1.966] | -0.284<br>[-0.569] | -0.649<br>[-1.561] | -0.038<br>[-0.055] |
| Exposure Earthquake (1/0) | -0.981<br>[-2.235] | -1.092<br>[-1.590] | -1.577<br>[-2.676] | -0.786<br>[-0.814] |
| Constant | 44.133<br>[136.824] | 153.771<br>[304.907] | 47.605<br>[115.279] | 165.589<br>[244.708] |
| Observations | 780 | 780 | 535 | 535 |
| R-squared | 0.022 | 0.058 | 0.039 | 0.026 |
| Log Likelihood | -4749 | -4749 | -3359 | -3359 |
| Notes : z-statistics in brackets |  |  |  |  |

### Models by severity of flu

Table 4 displays results from models for females estimated separately by flu severity in municipio of birth. These estimates are consistent with expectations as it is only among females born in high severity municipios that we find large effects of flu exposure on both adjusted height and knee height. Coefficients for knee height in high severity areas are at least twice as large as those in the first model. Effects on adjusted height are smaller but still noteworthy and in the expected direction.

**Table 4:**
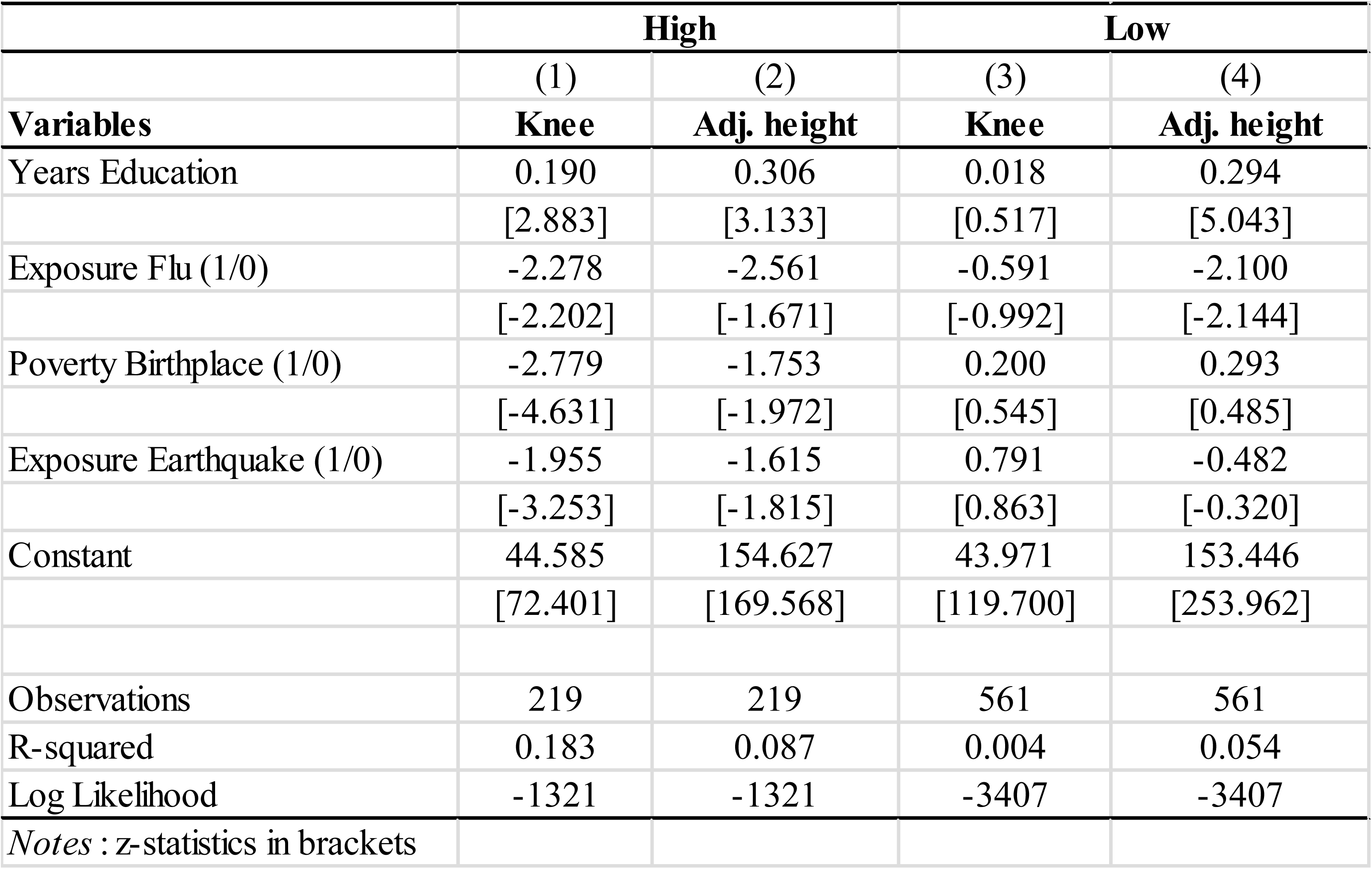
Models for females by flu severity of municipio of birth.

### A dangerous combo: the flu and the earthquake

Table 5 shows results of models estimated separately among those who were born in areas affected by the earthquake (high earthquake severity) and those born elsewhere (low earthquake severity). Again, consistent with expectations, the effects on knee height are stronger in areas hit by the earthquake. The magnitudes of effects on knee height among those born in Puerto Rico’s West coast are *three times larger* than those in previous models: knee reduction of those doubly exposed (e.g. to flu and earthquake) is equivalent to almost 3.5 cms, or about .82 of a standard deviation and .08 of mean knee height. The effects on adjusted height are noteworthy in both areas only of smaller relative magnitude (.4 of a standard deviation and .02 of the mean).

**Table 5:** Models for females by earthquake severity for females.

| Table 5: Models for females by earthquake severity for females |  |  |  |  |
| --- | --- | --- | --- | --- |
|  | High |  | Low |  |
|  | (1) | (2) | (3) | (4) |
| Variables | Knee | Adj. height | Knee | Adj. height |
| Years Education | 0.279 | 0.351 | 0.018 | 0.282 |
|  | [2.871] | [2.589] | [0.548] | [5.270] |
| Exposure Flu (1/0) | -3.459 | -3.292 | -0.716 | -2.137 |
|  | [-2.187] | [-1.490] | [-1.295] | [-2.400] |
| Poverty Birthplace (1/0) | -1.721 | -0.337 | -0.429 | -0.273 |
|  | [-1.922] | [-0.269] | [-1.271] | [-0.502] |
| Constant | 42.252 | 152.36 | 44.25 | 153.813 |
|  | [42.345] | [109.362] | [133.465] | [287.905] |
| Observations | 109 | 109 | 671 | 671 |
| R-squared | 0.159 | 0.085 | 0.006 | 0.052 |
| Log Likelihood | -657.3 | -657.3 | -4080 | -4080 |
| Notes : z-statistics in brackets |  |  |  |  |

These results suggest four inferences. First, the earthquake-tsunami and the flu pandemic had important effects but mostly among females. Second, the impacts of the flu are strong on knee height and in areas of high flu severity whereas those on height are less systematic. Third, and consistent with the idea that the combination of the two events was most consequential for the newly born and infants, the effects of flu exposure on knee height are two to three times larger among those who were simultaneously exposed to the earthquake. The flu effects in non-earthquake areas are in the proper direction but are of lesser magnitude. Given the definition of exposure we use here, these findings are very likely the result of perturbations of both the fetal *and* the postnatal period.

## DISCUSSION

There are two interpretations of our results that contradict our main conjecture. First, because our definition of exposure overlaps with the conventional definition, it could well be case that our estimates reflect impacts of *in utero* exposure, irrespective of postnatal experiences. To disprove this interpretation, we estimate models that include a dummy variable attaining the value 1 among those born in 1919. This is the definition used, among others, by Almond^(10)^ If only fetal exposure mattered, the estimated effects of the dummy should reflect it—as they do in previous studies-- and the coefficient of our preferred exposure variable should drift to zero. Table 6 displays results from a model estimated among female born in areas hit by the earthquake. The estimate of our preferred measure of exposure shows no changes whereas the dummy for birth year drifts to zero and/or is improperly signed. Similar inferences can be drawn from a model that uses quarter of birth (in 1919) as an index of fetal exposure.

**Table 6:** Model with conventional definition of exposure: females.

| **Table 6: Model with conventional definition of exposure: females** |  |  |
| --- | --- | --- |
|  | (1) | (2) |
| **Variables** | **Knee** | **Adj. height** |
| Years Education | 0.064 | 0.295 |
|  | [2.058] | [6.071] |
| Exposure Flu (1/0) | -1.368 | -2.085 |
|  | [-2.426] | [-2.371] |
| Year1919 (1/0) | 1.331 | -0.788 |
|  | [1.628] | [-0.618] |
| Constant | 43.630 | 153.487 |
|  | [161.404] | [364.141] |
| Observations | 780 | 780 |
| R-squared | 0.014 | 0.055 |
| LL | -4752 | -4752 |
| Notes : z-statistics in brackets |  |  |

Second, it is possible that our control variables (individual education and municipio’s poverty level) are insufficient to prevent contamination of estimates with the impact of conditions other than the flu and the earthquake. To partially remove this artifact we re-estimate the main models using municipios’ fixed effects. The new estimates of impacts are now net of municipios’ conditions correlated with exposure to the two events that have independent effects on nutritional status. Table 7 shows that the new estimates are *larger* than those in model with no fixed effects and that the associated p-values drop to less than .001.

**Table 7:** ESTIMATES IN MODELS WITH MUNICIPIOS FIXED EFFECTS: FEMALES.

| Table 7: ESTIMATES IN MODELS WITH MUNICIPIOS FIXED EFFECTS: FEMALES |  |  |  |  |
| --- | --- | --- | --- | --- |
|  | Not severe |  | Severe |  |
|  | (1) | (2) | (3) | (4) |
| Variables | Knee | Adj. height | Knee | Adj. height |
| yeduca | 0.468 | 0.248 | 0.324 | 0.474 |
|  | [1.395] | [4.305] | [3.541] | [3.567] |
| dummyF_1 | -0.778 | -3.935 | -3.980 | -2.720 |
|  | [-.446] | [-1.880] | [-2.744] | [-2.889] |
| Constant | 4.153 | 154.123 | 39.312 | 150.065 |
|  | [44.645] | [89.390] | [26.895] | [70.71] |
| Observations | 464 | 464 | 109 | 109 |
| R-squared | 0.017 | 0.023 | 0.327 | 0.209 |
| LL | -3763 | -3763 | -637 | 637 |
| Notes : z-statistics in brackets |  |  |  |  |
| Estimates of fixed effects omitted. Number of observations excludes cells with small number of observations |  |  |  |  |

### A placebo test

An alternative way to corroborate the causal role of the 1918 pandemic is to conduct a “placebo test” and show that the exposure indicator used here is unrelated to nutritional status markers among individuals *who were not exposed to the pandemic and/or the earthquake.* An ideal placebo test is impossible since, after all, the 1918 flu was a pandemic and virtually the entire world was exposed to it. However, our sample includes individuals who were *not born* in Puerto Rico and, therefore, were not exposed to the earthquake. If results are an artifact of unobserved variables, we should observe similar effects among the foreign born as we do among Puerto Rican natives in earthquake areas. However, a model estimated among the foreign born suggests that the flu effects in areas struck by the earthquake are close to zero. We hasten to emphasize that the test is underpowered because the foreign born constitute a small fraction of our sample (5 percent). However, our conjecture would have been crushed had we retrieved effects of the flu of even moderate size among the foreign born.

### Systematic errors, small sample sizes and the role of chance

Although estimates of effects on knee height are, by conventional standards, “statistically significant” (at levels p<.01 or less), we intentionally avoided use of this expression. Instead we prefer to refer to them as worthy of notice (or not). We did this for two reasons. The first is that the estimates could be contaminated by systematic measurement of errors. The second is that they may be due to chance.

#### Systematic errors

We conducted two tests to assess the possibility of systematic underestimation of knee height among those exposed to the flu *and* born in areas impacted by the earthquake. First, the distribution of knee height shows no deviant extreme cases and the smallest values in earthquake areas are within 1.5 of a standard deviation. Second, in a more radical test we *ignored* the lowest values of knee height and re-estimated models. The key inference from this exercise is the following: to convert estimated effects from “worthy of notice” (p < [.01-.02]) to mundane (p>.02) we *need to exclude observations of knee height below the first quartile* of the distribution, a rather radical and also implausible surgery.

#### The role of chance

Most epidemiological and population health research highlights findings on the basis of classic-Fisher criteria, that is, based on *a priori* chosen significance level (say α<.025 in a two-tailed test). We are saying nothing knew when we point out that this type of criterion can be highly misleading. This is of particular concern as extreme values of a statistic can be obtained just by chance. To assess this possibility, we pursue two routes: (a) perform a permutation test ^(48)^ and (b) compute bounds for false discovery rates ^(49)^.

##### (a) Permutation test

we implement the simplest of permutation tests and verify that it leads to the same inferences drawn from conventional hypothesis testing, namely, effects of knee height of the size we observe occur with probabilities lower than [.01-.02] (see Figure in S4 Text)

##### (b) False discovery rate

this is the *conditional* probability that if the null hypothesis is rejected, it is erroneously rejected. This quantity is usually quite different from the conventional α as it is a function of α, power, and the true magnitude of effects. Alternative values of these parameters produce results (see figure in S5 figure) that confirm our inferences. Indeed, given our p-values ([.01-.02]) and approximate power (.50-.60), the probability of uncovering effects only by chance is between .10 and .30, hardly a comforting range but quite common in clinical and population studies^(51)^.

#### How large are the estimated effects?

Is the size of the 1918 flu effects on survivors nutritional status significant? While our empirical findings confirm that flu exposure, in the broadest sense defined here, is robustly associated with markers of early nutritional status, it is unclear whether the magnitude of effects is substantively meaningful. To provide a sense of magnitude we compare predicted changes in individual stature associated with flu exposure with changes in stature throughout the period of mortality decline in Western Europe. Since there are no historical records of knee height, we exploit the fact that knee height is strongly associated with height and draw tentative inferences after predicting changes in height using estimated changes in knee height. Although the observed association between height and knee height in our sample is contaminated by systematic errors in height due to skeletal compression, it is in all likelihood underestimated. Thus, the estimated slope of the regression of adjusted height on knee height is biased downwards and predicted values of height given knee height will be underestimated.

A log-log regression of adjusted height on knee height reveals that the reduction in height implied by the reduction in knee height due to flu and earthquake exposure estimated before (in the range 1.5-6.5 cms) is associated with a proportionate adjusted height reduction of about .0243. To place this in context, consider this: it took forty years, between 1860 and 1900, for the mean height of the Dutch population to experience proportionate gains of about .012!.

A final piece of empirical evidence boosts the significance of the effects we find in the data. We use information on female respondents’ inter-wave mortality and estimate a model including as predictor the variable knee-height. The effect of this variable is powerful as an increase of 1 cm in knee height translates into a *decrease* in mortality risks above age 75 of the order of 8 %. A decrease of this magnitude in life tables for the US during the period 2000-2010 (the period of time covered by the PREHCO survey) is equivalent to an increase in female life expectancy at age 75 from 12 to 13 years over a period of 20 years. Since the reduction effect on knee height due to the flu and earthquake combined is within the range (1.5-6.5 cms), the implication is that survivors of these cohorts of females might have lost between .4 and 2.5 years of residual life expectancy or, equivalently, between 40 and 250 percent of the gains experienced in a period of 20 years. These are not trivial effects.

## CONCLUSION

We argued that past research on the long-run effects of the 1918 influenza may be blindsided by a preoccupation with fetal exposure. Although there is strong evidence supporting the idea that embryonic and other intrauterine disruptions are influential, fetal development is also about growth of cartilage, bone and muscle tissue, all of which are implicated *in subsequent postnatal physical development*. Furthermore, impairments in the fetal period can be aggravated if post-natal conditions are also unfavorable. This justifies our claim that the flu pandemic could have also perturbed the post-natal period and through both, fetal and postnatal exposures, affected children’s nutritional status. Our estimates, particularly those for female in born in high severity areas and/or in earthquake-tsunami zones, are large, statistically “relevant”, and robust to checks. This evidence does not imply that fetal exposure is irrelevant but that it, together with postnatal conditions, combine in a highly poisonous cocktail that impedes attainment of physical growth landmarks.

The paper has shortcomings. First, only one of two markers of nutritional status, knee height, is systematically responsive to flu/earthquake exposure. This could be due to the fact that adjusted height is more likely to be influenced by measurement errors than knee height. As a result, it is difficult to tell whether the unequal response could also be due to differences in physiological processes that underpin development of different parts of the human body. Second, the sample is small and vulnerable to produce effects where there are none. Unlike other research, we are not dealing with observations in the tens of thousands but with an effective sample size orders of magnitude below that. This does not favor strong model fit even though goodness of fit is always strongest in models with flu/earthquake exposure indicators. However, despite the noise, there is a strong and systematic signal that resists multiple checks. Admittedly, these checks can only suggest and will never prove that results are immune to false discovery and other aberrations produced by the data or by chance. Finally, a word about the target population. Puerto Rico is a tiny dot in a world map. Its population size has always been, then and now, an infinitesimal fraction of the world population. Why would anybody bother with all of this? First, the unlikely collusion of two simultaneous natural disasters and the accidental availability of empirical records of survivors, generated a unique opportunity to identify effects of broadly defined early exposures to shocks. Second, we find stubborn empirical evidence suggesting that perhaps past research on the impacts of the 1918 flu pandemic may have missed something important: the influence of the combined disruption of fetal and postnatal life on the ultimate fate of subsequent physical growth. We are not so much trumpeting a new finding as we are identifying a relation that deserves a second look in future research with pandemics of similar nature.

## ACKNOWLEDGEMENTS

We thank Berty Lumey for helpful comments to the first draft of the manuscript.

## SUPPORTING MATERIAL CAPTIONS

S1 Text: **FLU SEVERITY**

S2 Text : **ADJUSTED HEIGHT**

S3 Table: **MODELS FOR MALES**

S4 Text: **RESULTS OF PERMUTATION TEST**

S5 Figure: ALTERNATIVE RATES OF FALSE DISCOVERY

